# Summer dynamics of microbial diversity on a mountain glacier

**DOI:** 10.1101/2022.06.04.494832

**Authors:** Scott Hotaling, Taylor L. Price, Trinity L. Hamilton

## Abstract

Under climate change, glaciers are rapidly receding worldwide. A melting cryosphere will dramatically alter global sea levels, carbon cycling, and water resource availability. Glaciers also host rich biotic communities that are dominated by microbial diversity and this biodiversity can impact surface albedo, thereby driving a feedback loop between biodiversity and cryosphere melt. However, the microbial diversity of glacier ecosystems remains largely unknown outside of major ice sheets, particularly from a temporal perspective. Here, we characterized temporal dynamics of bacteria, eukaryotes, and algae on the Paradise Glacier, Mount Rainier, USA, over the summer melt season. During our study, the glacier surface steadily darkened as seasonal snow melted and darkening agents accumulated until new snow fell in late September. From a community-wide perspective, the bacterial community remained generally constant and eukaryotes exhibited a clear temporal progression of community change while fungal diversity was intermediate. Individual taxonomic groups, however, exhibited considerable stochasticity. We found little support for our *a priori* prediction that autotroph abundance would peak before heterotrophs. Notably, two different trends in snow algae emerged—an abundant early-and late-season OTU with a different mid-summer OTU that peaked in August. Overall, our results highlight the need for temporal sampling to clarify microbial diversity on glaciers and that caution should be exercised when interpreting results from single or few timepoints.

## Body

Glacier ecosystems are key components of global biodiversity and support diverse, mostly microbial communities comprised of bacteria, photosynthetic algae, and fungi [8, 9, 15].However, beyond point estimates of biodiversity, seasonal variation of these biota is poorly understood. To date, the majority of biological research on glaciers has focused on establishing baselines of biodiversity [4], understanding the ecophysiology of resident organisms [3], resource availability and use [7, 13], and clarifying drivers of biological albedo reduction [where pigmented organisms darken the cryosphere and promote melt, 9]. However, temporal perspectives of biodiversity in glacier ecosystems remain rare [but see 5, 14, 16].

Glacier surfaces are highly dynamic and experience substantial environmental fluxes in space and time. Early season “spring” conditions on temperate glaciers are typically marked by increasing periods of daylight with intense temperature swings and relatively little biological activity. By summer, temperature swings have moderated and biotic activity including photosynthesis, respiration and nutrient cycling near annual peaks [1]. In fall, days shorten, temperatures decrease, and snowfall events limit primary productivity [1].

Here, we present a temporal perspective of microbial community change on the Paradise Glacier, Mount Rainier, WA, USA (Fig. 1a,b), a temperate alpine glacier that hosts a diverse, representative community of glacier biota. From May to September, 2019, we collected triplicate snow samples from ∼2255 m on the eastern margin of the glacier and tracked changes in microbial communities by sequencing 16S and 18S small subunit rRNA and fungal ITS amplicons (detailed methods provided in Supporting Information). We expected to uncover a rich biological community on the glacier and evidence of successional dynamics with primary producer abundance peaking early in summer followed by an increase in heterotrophs later in the season.

**Figure 1.**
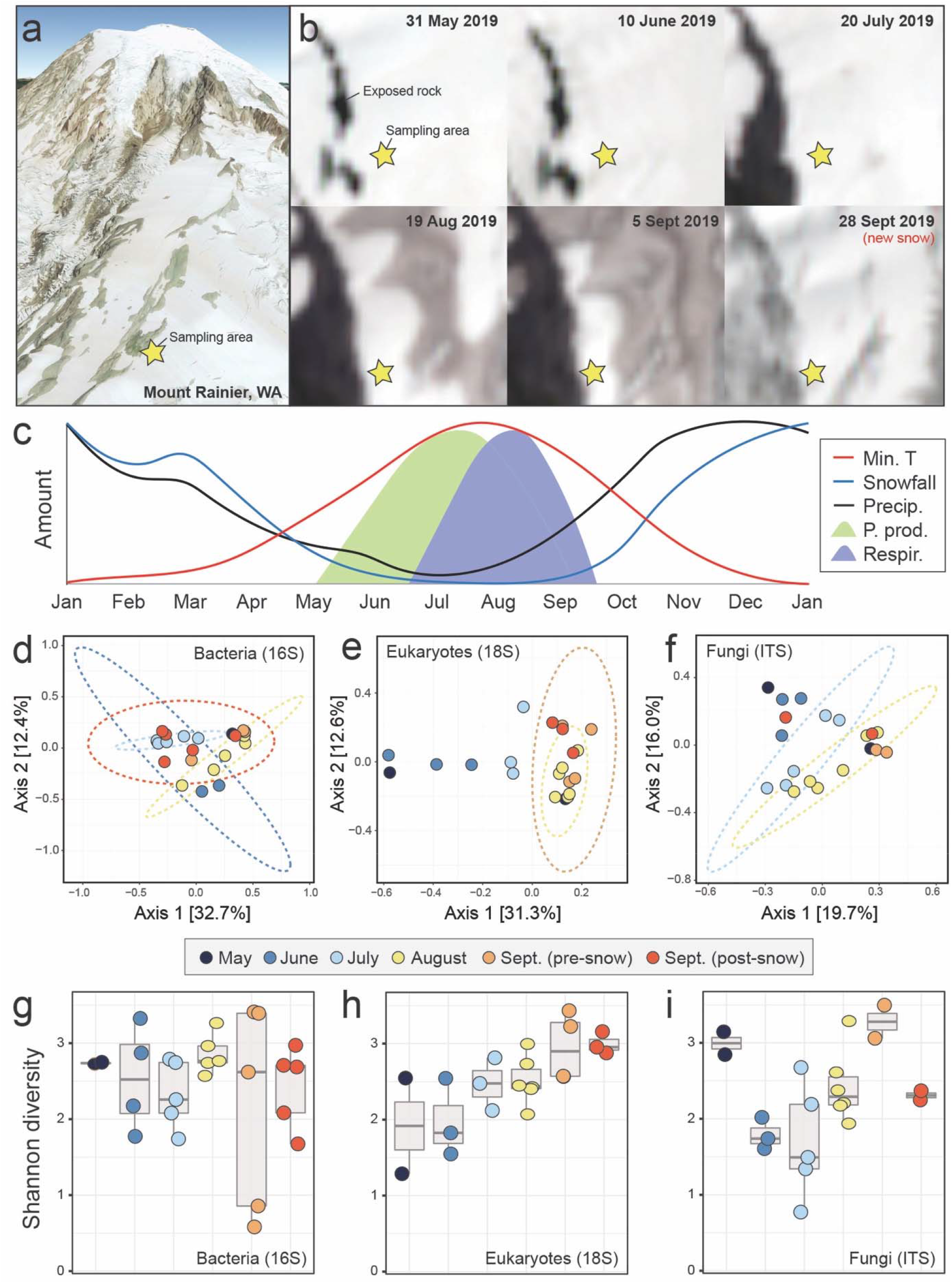
(a) Location of our study site on the Paradise Glacier of Mount Rainier, Washington, USA. Imagery from Google Earth. (b) Sentinel-2 satellite imagery of the study site from late May to September, 2019. A fresh snowfall occurred between the final two sampling timepoints in September. (c) A conceptual image of primary production and heterotroph activity on a temperature mountain glacier over the course of one year. Overlaid on this conceptual framework are monthly averages of minimum temperature, average precipitation, and average snowfall for the nearby Paradise Ranger Station (1655 m) from 1916-2016 (data from the Western Regional Climate Center). (d-f) Principal coordinate analysis of community composition based on Bray-Curtis dissimilarity for (d) bacteria, (e) eukaryotes, (f) and fungi. (g-i) Shannon diversity through time for the same sampling points and communities: (g) bacteria, (h) eukaryotes, and (i) fungi.

During our study, the Paradise Glacier surface darkened as seasonal snow receded, debris accumulated, and biotic processes (e.g., snow algal blooms) transpired until late September when new snow fell (Fig. 1a-b). Overall, we recovered 4724 bacterial OTUs (16S), 1246 eukaryotic OTUs (18S), and 3007 fungal OTUs (ITS). The bacterial community was distinct month-to-month, particularly later in the season (Fig. 1d). The eukaryotic and fungal communities were less clearly differentiated month-to-month but exhibited more seasonal progression than bacteria (i.e., the amount of time between sampling events appeared to generally scale with community turnover; Figs. 1e,f). Alpha diversity (Shannon’s) was temporally stable for bacteria (Fig. 1g), steadily increased for eukaryotes (Fig. 1h), and was variable for fungi (Fig. 1i). The effects of September snowfall had little effect on the community composition, alpha diversity, nor relative abundances (Figs. 1d-i, 2).

The most abundant bacterial OTUs were affiliated with Bacteroidetes and Proteobacteria (Fig. 2a-b). Within the Bacteroidetes, OTUs assigned to *Ferruginibacter* and *Solitalea* were most abundant and OTUs assigned to *Pseudomonas* (Gammaproteobacteria) and *Exiguobacterium* (Bacillota) were also common. For eukaryotes, OTUs assigned to green algae were abundant, including four Chlorophyta OTUs; three were assigned to the snow algae genus *Chlainomonas* while the fourth belonged to Cyanidiales. Basidiomycota OTUs were prevalent in the fungal data, including six of the 10 most abundant OTUs. These six OTUs were affiliated with the Microbotryomycetes including *Phenoliferia* and *Filobasidium* as well as OTUs that could not be classified below the Class level.

**Figure 2.**
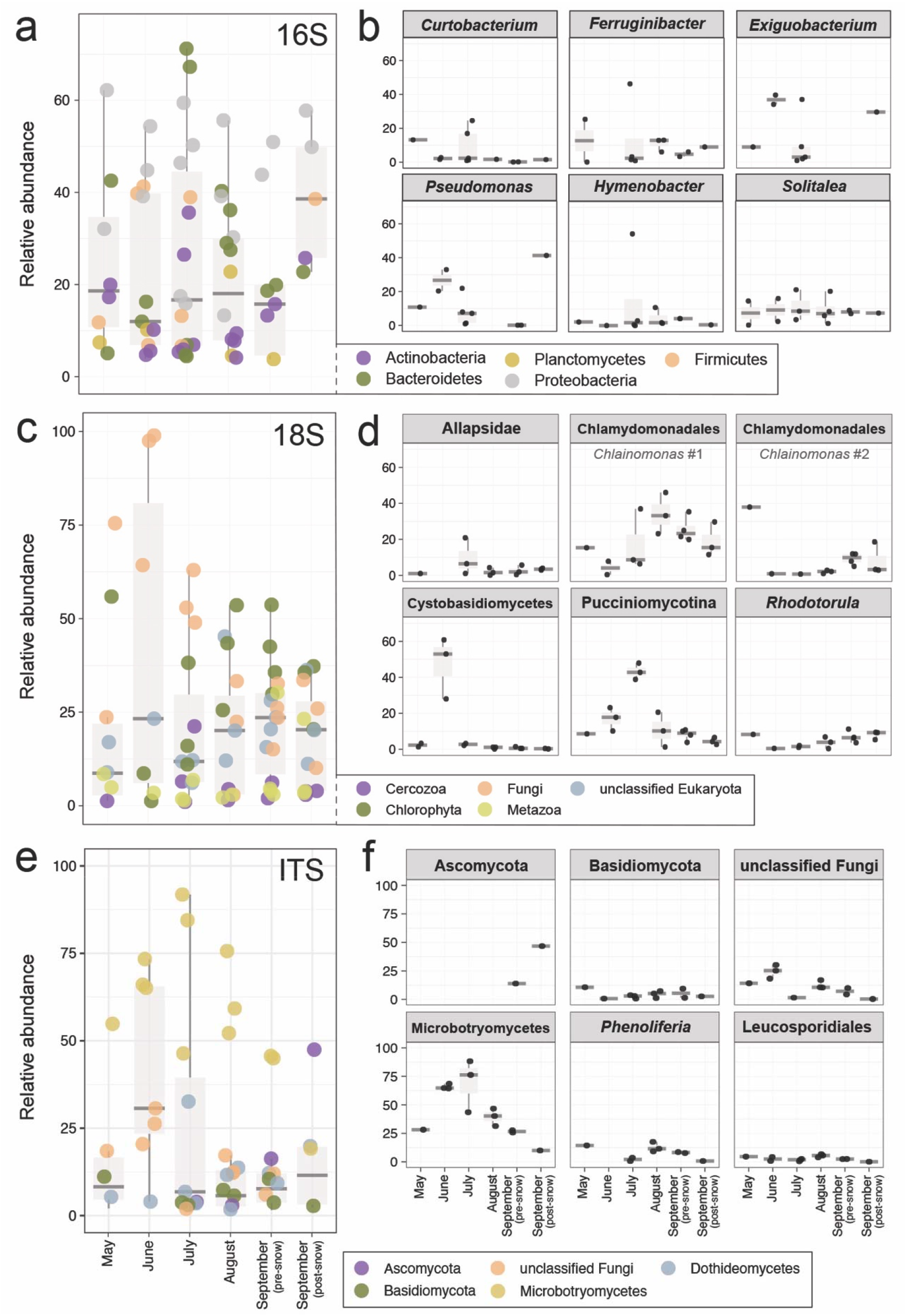
Temporal abundance of common taxonomic groups for each data set overall and broken down for select taxa: (a-b) 16S rRNA, (c-d) 18S rRNA, and (e-f) fungal ITS. Taxonomic groups comprising the largest percent relative abundance in each library are shown in a, c, and e. The most abundant operational taxonomic units (OTUs) in each data set are shown in b, d and f where taxonomy has been assigned to each OTU at the highest resolution possible (see detailed methods in the Supporting Information). Box plots show mean percent relative abundance of the group (a, c and e) or OTU (b, d and f). Data sets are binned by month of sample collection except for early and late September.

The abundance of most major bacterial groups fluctuated through time (e.g., Bacteroidetes and Actinobacteria were most abundant in July and less abundant in early September, Fig. 2a). In contrast, Proteobacteria were abundant in all samples. Algal taxa (phylum Chlorophyta), perhaps the most influential eukaryotes on glaciers [9], were recovered in all samples from all months (Fig. 2c) but were least abundant in June. Algal community composition shifted throughout the summer: abundant *Chlainomonas* OTUs in May and late September were distinct from those recovered in July-September samples (Fig. 2d). For fungi, the relative abundance of sac fungi (Ascomycota) increased in late summer, peaking after the first significant snowfall in September (Fig. 2e). Conversely, the highest abundances of Basidiomycota (the other division that comprises the subkingdom Dikarya alongside Ascomycota) were observed in May with lower levels from June-September (Fig. 2f).

Broadly, our results support dynamism in both taxonomic composition and abundance of microbial communities on mountain glaciers during the summer melt season. For many groups (e.g., *Ferruginibacter*, Fig. 2b), abundance trends appeared stochastic, or at least not linked to any seasonal dynamics, while others (e.g., Pucciniomycota) exhibited clear directionality across the melt season. Given the resource-poor nature of glacier ecosystems [13], we expected to observe an early-season wave of primary producers followed by an increase in heterotrophs later in the season. Contrary to our expectation, OTUs for snow algal primary producers, particularly *Chlainomonas* within the Chlorophyta, were abundant in all samples except June.

Because the same May *Chlainomonas* OTUs increased in abundance in late September, May samples could reflect cells buried from previous years. In contrast, the July—September samples contain *Chlainomonas* OTUs that are distinct from this “resident” community and are perhaps the product of atmospheric input. However, since physical and chemical snow conditions can impact snow algae composition and pigment content [12] which vary seasonally [11], it is also possible that both algal communities are present and local conditions drive the differences we observed. We did observe a decrease in Microbotryomycetes (in the Basidiomycota) and an increase in Ascomycota, fungi which typically favor nutrient-rich niche space [2], in later season samples. Shifts in fungal taxa in response to temperature and nutrients [10] have been linked to resource availability selecting for specific taxa.

With widespread interest in microbial diversity in the cryosphere to better understand carbon cycling, biological albedo reduction, and community ecology of glacier ecosystems [1, 6, 13], it is clear that one or a few estimates of abundance may not reflect broader trends. Thus, our data underscore the need for temporal sampling to ultimately uncover higher level links between biology and the cryosphere in the mountain cryosphere [9]. To realize this potential, such efforts should ideally occur across multiple locations within and among montane regions.

## Acknowledgements

T.L.H. was supported by NSF awards #EAR-1904159 and #EAR-2113784. Samples were collected under research permit #MORA-2018-SCI-0005. The authors acknowledge the Minnesota Supercomputing Institute at the University of Minnesota for providing resources that contributed to the research results reported within this paper.

## Author contributions

S.H. and T.L.H. conceived of the study. S.H. collected samples. T.L.H. performed analyses.

S.H. and T.L.H. wrote the manuscript with support from T.P. All authors read and approved the final version.

## References

1. Anesio AM, Laybourn-Parry J (2012). Glaciers and ice sheets as a biome. Trends Ecol Evol 27;219–225.

2. Crowther TW, Boddy L T Jones H (2012). Functional and ecological consequences of saprotrophic fungus–grazer interactions. The ISME Journal 6;1992–2001.

3. Dial RJ, Becker M, Hope AG, Dial CR, Thomas J, Slobodenko KA et al (2016). The role of temperature in the distribution of the glacier ice worm, Mesenchytraeus solifugus (Annelida: Oligochaeta: Enchytraeidae). Arctic, Antarctic, and Alpine Research 48;199–211.

4. Edwards A, Pachebat JA, Swain M, Hegarty M, Hodson AJ, Irvine-Fynn TDL et al (2013). A metagenomic snapshot of taxonomic and functional diversity in an alpine glacier cryoconite ecosystem. Environmental Research Letters 8;035003.

5. Els N, Greilinger M, Reisecker M, Tignat-Perrier R, Baumann-Stanzer K, Kasper-Giebl A et al (2020). Comparison of bacterial and fungal composition and their chemical interaction in free tropospheric air and snow over an entire winter season at Mount Sonnblick, Austria. Frontiers in microbiology 11;980.

6. Ganey GQ, Loso MG, Burgess AB, Dial RJ (2017). The role of microbes in snowmelt and radiative forcing on an Alaskan icefield. Nature Geoscience 10;754.

7. Hamilton TL, Havig JR (2018). Inorganic carbon addition stimulates snow algae primary productivity. The ISME journal;1.

8. Hotaling S, Hood E, Hamilton TL (2017). Microbial ecology of mountain glacier ecosystems: biodiversity, ecological connections and implications of a warming climate. Environmental microbiology 19;2935–2948.

9. Hotaling S, Lutz S, Dial RJ, Anesio AM, Benning LG, Fountain AG et al (2021). Biological albedo reduction on ice sheets, glaciers, and snowfields. Earth-Science Reviews 220;103728.

10. Li H, Yang S, Semenov MV, Yao F, Ye J, Bu R et al (2021). Temperature sensitivity of SOM decomposition is linked with a K-selected microbial community. Global Change Biology 27;2763–2779.

11. Onuma Y, Takeuchi N, Tanaka S, Nagatsuka N, Niwano M, Aoki T (2020). Physically based model of the contribution of red snow algal cells to temporal changes in albedo in northwest Greenland. The Cryosphere 14;2087–2101.

12. Remias D, Karsten U, Lütz C, Leya T (2010). Physiological and morphological processes in the Alpine snow alga Chloromonas nivalis (Chlorophyceae) during cyst formation. Protoplasma 243;73–86.

13. Ren Z, Martyniuk N, Oleksy IA, Swain A, Hotaling S (2019). Ecological stoichiometry of the mountain cryosphere. Frontiers in Ecology and Evolution 7;360.

14. Sheik CS, Stevenson EI, Den Uyl PA, Arendt CA, Aciego SM, Dick GJ (2015). Microbial communities of the Lemon Creek Glacier show subtle structural variation yet stable phylogenetic composition over space and time. Frontiers in microbiology 6;495.

15. Stibal M, Bradley JA, Edwards A, Hotaling S, Zawierucha K, Rosvold J et al (2020). Glacial ecosystems are essential to understanding biodiversity responses to glacier retreat. Nature Ecology & Evolution 4;686–687.

16. Winkel M, Trivedi CB, Mourot R, Bradley JA, Vieth-Hillebrand A, Benning LG (2022). Seasonality of Glacial Snow and Ice Microbial Communities. Frontiers in microbiology;1760.

