## Supporting Information for "Summer dynamics of microbial diversity on a mountain glacier"

**METHODS**

**Sample collection and DNA extraction**

Samples were collected on Paradise Glacier in May, June, July, August, and September 2019.

For each sample, the uppermost ~1 cm of the glacier was scraped with an ethanol-sterilized collection spoon into a 500 mL Whirl-Pak bag until it was filled. Samples were transported on ice and remained frozen until processing. In the lab, Total genomic DNA was extracted using a DNeasy PowerSoil Kit (Qiagen, Carlsbad, CA, USA). The concentration of DNA was determined using a Qubit RNA Assay kit (Molecular Probes, Eugene, OR, USA) and a Qubit 3.0 Fluorometer (Life Technologies, Carlsbad, CA).

**Amplicon sequencing**

Total DNA for each sample was submitted to the University of Minnesota Genomics Center (UMGC) for amplicon sequencing. Amplicons were sequenced at UMGC using MiSeq Illumina 2 × 300 bp chemistry with the primers 515Ff and 806rB targeting the V4 region of bacterial and archaeal 16S SSU rRNA gene sequences (Caporaso et al., 2012; Apprill et al., 2015), primers E572F and E1009 targeting the V9 region of eukaryotic 18S SSU rRNA gene sequences (Comeau et al., 2011), and primers ITS1F and ITS2R for ITS1 (Tedersoo et al., 2015). UMGC prepared dual indexed Nextera XT DNA libraries following their improved protocol for library preparation which enables detection of taxonomic groups that often go undetected with existing methods (Gohl et al 2016). Each sample was sequenced once.

**Amplicon analysis**

Post sequence processing was performed using the mothur (ver. 1.45.3) sequence analysis platform (Schloss et al., 2009) following the MiSeq SOP (Kozich et al., 2013) as described previously (Havig and Hamilton, 2019). For 16S and 18S rRNA, read pairs were assembled and resulting contigs with ambiguous bases were removed and trimmed to include only the overlapping regions. For ITS amplicons, we proceeded with only the forward reads by removing ambiguous bases. Chimeras were identified and removed using UCHIME (Edgar et al., 2011) from all the data sets. 16S rRNA amplicons were aligned against the SILVA v138 database and clustered into operational taxonomic units (OTUs) at a sequence similarity of 0.97 with the OptiClust algorithm in mothur. 16S rRNA OTUs classified within mothur using the SILVA database (v138). 18S rRNA amplicons were aligned against the PR^2^ database (ver. 4.12.0) (Guillou et al., 2013).18S rRNA OTUs clustered into OTUs at a sequence similarity of 0.99 using the OptiClust algorithm in mothur. 18S rRNA OTUs and classified within mothur using the PR2 database (v4.12.0). ITS OTUs were clustered into OTUs at a sequence similarity 0.97 using the agc algorithm in mothur. ITS OTUs were classified within Mothur using the UNITE database (v8.2) (Abarenkov et al., 2020).

**Statistical analysis**

All post-mothur processing was carried out in R (ver. 4.0.3; R Core Team, 2018). We calculated diversity indices using vegan (Oksanen et al., 2017) within the Phyloseq package (McMuride and Holnes, 2013) and data were visualized using ggplot2 (Wickham 2016). Data were rarefied to the lowest sequence coverage by random subsampling for richness calculations and transformed using a variance stabilized transformation (Anders and Huber 2010). Diversity was calculated using the Bray-Curtis dissimilarity metric on the transformed data.

**Data availability**

All raw sequences are available through the NCBI Sequence Read Archive under BioProject number PRJNA799302.

**References**

Abarenkov K., Zirk A., Piirmann T., Pöhönen R., Ivanov F., Nilsson, Henrik R.,; Kõljalg U. (2020) UNITE mothur release for Fungi. Version 04.02.2020. UNITE Community. https://doi.org/10.15156/BIO/786381

Edgar R. C., Haas B. J., Clemente J. C., Quince C., and Knight R. (2011) UCHIME improves sensitivity and speed of chimera detection. *Bioinformatics* 27, 2194–2200. doi: 10.1093/bioinformatics/btr381

Gohl D. M., Vangay P., Garbe J., MacLean A., Hauge A., Becker A., et al. (2016) Systematic improvement of amplicon marker gene methods for increased accuracy in microbiome studies. *Nature* 201:6.

Guillou L., Bachar D., Audic S., Bass D., Berney C., Bittner L., et al. (2013) The Protist Ribosomal Reference database (PR2): A catalog of unicellular eukaryote small sub‐unit rRNA sequences with curated taxonomy. *Nucleic Acids Research*, 41, D597–D604.

Havig, J. R. & Hamilton, T.L. (2019) Cryptic oxygen oases: Hypolithic photosynthesis in hydrothermal areas and implications for Archean surface oxidation. *Front. Earth Sci*. 7:15.

Kozich J. J., Westcott S. L., Baxter N. T., Highlander S. K., Schloss P. D. (2013) Development of a dual-index sequencing strategy and curation pipeline for analyzing amplicon sequence data on the MiSeq Illumina sequencing platform. *Appl. Environ. Microbiol*. 79, 5112–5120. doi: 10.1128/AEM.01043-13

McMurdie P. J. & Holmes S. (2013) phyloseq: an R package for reproducible interactive analysis and graphics of microbiome census data. *PloS One*, 8(4), e61217.

Oksanen J., Blanchet F.G., Friendly M., Kindt R., Legendre P., McGlinn D., et al. (2017) vegan: community ecology package. R package ver- sion 2.4-2. https://cran.r-project.org/web/packages/vegan/index.html.

R Core Team (2018) R: a language and environment for statistical computing. R Foundation for Statistical Computing. https://www.r-project.org/

Schloss P.D., Westcott S.L., Ryabin T., Hall J.R., Hartmann M., Hollister E.B., Lesniewski R.A., Oakley B.B., Parks D.H., Robinson C.J. and Sahl J.W. (2009) Introducing mothur: open-source, platform-independent, community-supported software for describing and comparing microbial communities. *Appl. Environ. Microbiol*., 75(23), 7537-7541.

Tedersoo L., Anslan S., Bahram M., Põlme S., Riit T., Liiv I., et al. (2015) Shotgun met- agenomes and multiple primer pair-barcode combinations of ampli- cons reveal biases in metabarcoding analyses of fungi. *MycoKeys* 10: 1–43.

Wickham H. (2016) ggplot2: Elegant Graphics for Data Analysis. Springer-Verlag New York ISBN 978-3-319-24277-4
